# Complexity of the eukaryotic dolichol-linked oligosaccharide scramblase suggested by activity correlation profiling mass spectrometry

**DOI:** 10.1101/2020.04.15.043679

**Authors:** Alice Verchère, Andrew Cowton, Aurelio Jenni, Monika Rauch, Robert Häner, Johannes Graumann, Peter Bütikofer, Anant K. Menon

## Abstract

The canonical pathway of *N*-linked protein glycosylation in yeast and humans involves transfer of the oligosaccharide moiety from the glycolipid Glc_3_Man_9_GlcNAc_2_-PP-dolichol to select asparagine residues in proteins that have been translocated into the lumen of the endoplasmic reticulum (ER). Synthesis of Glc_3_Man_9_GlcNAc_2_-PP-dolichol occurs in two stages, producing first the key intermediate Man_5_GlcNAc_2_-PP-dolichol (M5-DLO) on the cytoplasmic face of the ER, followed by translocation of M5-DLO across the ER membrane to the luminal leaflet where the remaining glycosyltransfer reactions occur to complete the structure. Despite its critical importance for *N*-glycosylation, the scramblase protein that mediates the translocation of M5-DLO across the ER membrane has not been identified. Building on our ability to recapitulate scramblase activity in large unilamellar proteoliposomes reconstituted with a crude mixture of ER membrane proteins, we developed a mass spectrometry-based ‘activity correlation profiling’ approach to identify scramblase candidates in the yeast *Saccharomyces cerevisiae*. Curation of the activity correlation profiling data prioritized six polytopic ER membrane proteins as scramblase candidates, but reconstitution-based assays and gene disruption in the protist *Trypanosoma brucei* revealed, unexpectedly, that none of these proteins were necessary for M5-DLO scramblase activity. Our results instead suggest the possibility that the M5-DLO scramblase may be a protein, or protein complex, whose activity is regulated at the level of quaternary structure. This key insight will aid future attempts to identify the scramblase.

## Introduction

Asparagine *(N)*-linked protein glycosylation is found in all domains of life (Breitling, J. and Aebi, M. 2013, Hirschberg, C.B. and Snider, M.D. 1987, Larkin, A. and Imperiali, B. 2011, Lennarz, W.J. 1987, Schenk, B., Fernandez, F., et al. 2001). In eukaryotes, it takes place in the lumen of the endoplasmic reticulum (ER), when oligosaccharyltransferase (OST) attaches a pre-synthesized oligosaccharide (Glc_3_Man_9_GlcNAc_2_ in yeast and humans) to asparagine residues within glycosylation sequons in newly translocated proteins (Kelleher, D.J. and Gilmore, R. 2006, Shrimal, S. and Gilmore, R. 2019). The Glc_3_Man_9_GlcNAc_2_ oligosaccharide is built by sequentially adding sugars to dolichyl phosphate, an isoprenoid glycosyl carrier lipid with a very long hydrocarbon chain (Fig. 1A). Sugar addition occurs in two stages and on different sides of the ER membrane. The first stage comprises seven reactions in which the sugar nucleotide donors UDP-GlcNAc and GDP-Man are used to produce the key intermediate Man_5_GlcNAc_2_-PP-dolichol (M5-DLO) on the cytoplasmic face of the ER (Fig. 1A). M5-DLO is then translocated to the luminal side of the ER where seven, lipid-mediated glycosyltransfer steps occur to generate Glc_3_Man_9_GlcNAc_2_-PP-dolichol (G3M9-DLO). The mannose and glucose residues needed for these latter reactions are donated by mannose phosphate dolichol (MPD) and glucose phosphate dolichol (GPD), both of which are synthesized on the cytoplasmic face of the ER and flipped to the luminal leaflet (Anand, M., Rush, J.S., et al. 2001, Haselbeck, A. and Tanner, W. 1982, Rush, J.S., van Leyen, K., et al. 1998, Rush, J.S. and Waechter, C.J. 1995, Sanyal, S. and Menon, A.K. 2010, Schenk, B., Imbach, T., et al. 2001, Skrabanek, L.A. and Menon, A.K. 2018).

**Figure 1:**
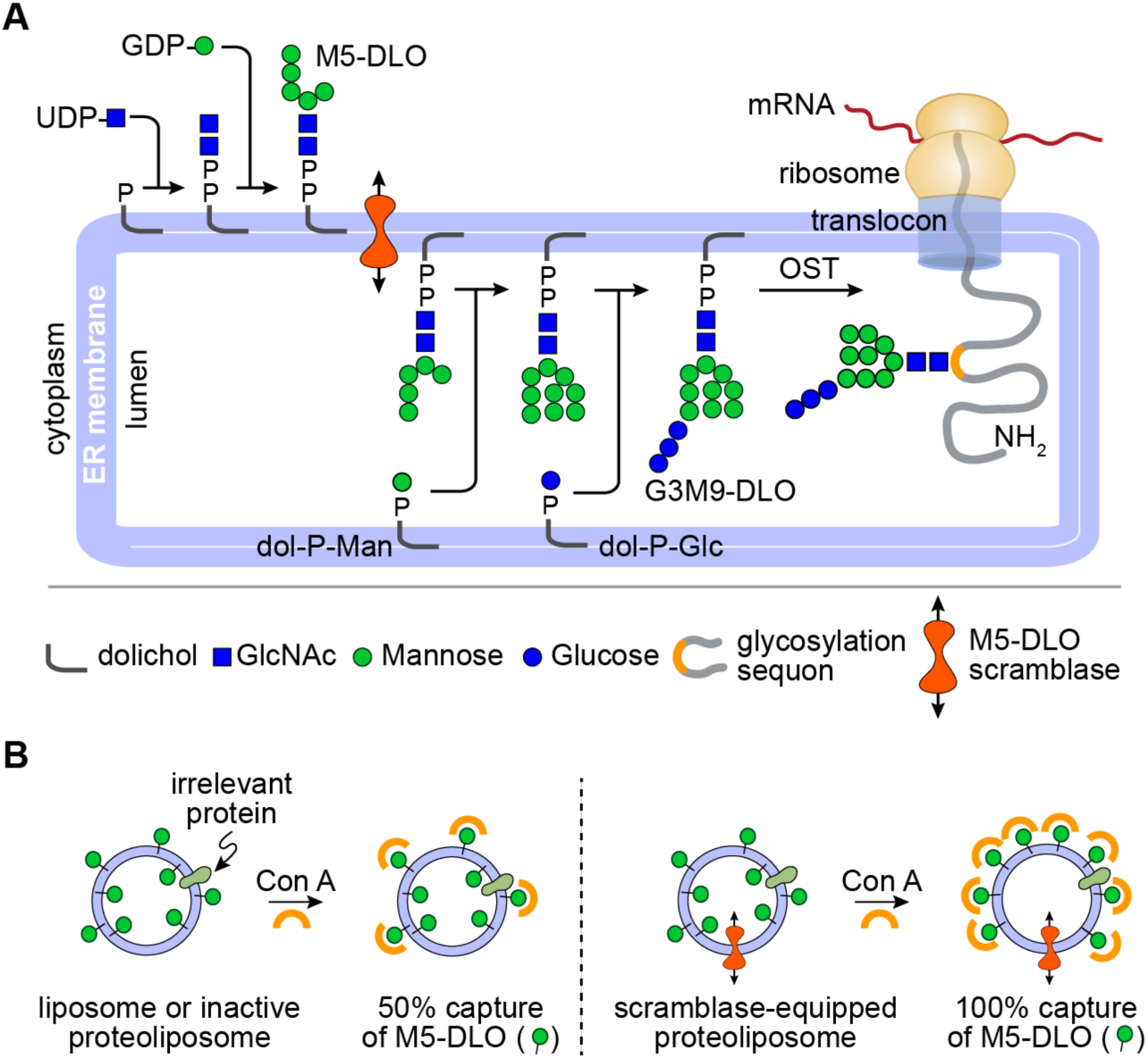
Transbilayer movement of Man_5_GlcNAc_2_-PP-dolichol (M5-DLO). **A**. Synthesis of Glc_3_Man_9_GlcNAc_2_-PP-dolichol (G3M9-DLO), the oligosaccharide donor for protein *N*-glycosylation in yeast and humans, occurs in two stages. The first stage, comprising seven glycosyltransfer reactions, occurs on the cytoplasmic side of the ER to generate M5-DLO from dolichyl phosphate, UDP-GlcNAc and GDP-mannose. The remaining seven glycosyltransfer reactions occur in the ER lumen, adding four mannose residues (from dolichylphosphoryl mannose (dol-P-man)) and three glucose residues (from dolichylphosphoryl glucose (dol-P-Glc)) to M5-DLO to generate G3M9-DLO. The switch from the cytoplasmic to the lumenal side of the ER requires transbilayer movement of M5-DLO, which is facilitated by M5-DLO scramblase (red icon). The right side of the figure shows oligosaccharyl-transferase (OST)-mediated transfer of the Glc_3_Man_9_GlcNAc_2_ oligosaccharide from G3M9-DLO to an asparagine residue within a glycosylation sequon in a nascent protein as it emerges from the protein translocon into the ER lumen. See text for details. **B**. Recapitulation of M5-DLO scrambling in a reconstituted system. The assay exploits the organic-solvent-resistant interaction between M5-DLO and the lectin Concanavalin A (Con A) - thus whereas M5-DLO is soluble in organic solvent, M5-DLO:ConA complexes are precipitated. Proteoliposomes are reconstituted from egg phosphatidylcholine, [^3^H]M5-DLO and ER membrane proteins. Protein-free liposomes are prepared in parallel. Incubation of the liposomes with Con A results in the capture/precipitation of ∼50% of [^3^H]M5-DLO (left), corresponding to the pool in the outer leaflet of the vesicles, whereas similar incubation of proteoliposomes containing M5-DLO scramblase results in the capture of ∼100% of [^3^H]M5-DLO (right) as M5-DLO molecules in the inner leaflet are transported to the outer leaflet where they are captured by Con A. Proteoliposomes reconstituted with proteins other than the M5-DLO scramblase, i.e. irrelevant proteins, are expected to behave similarly to protein-free liposomes (left).

The assembly of G3M9-DLO thus requires the rapid translocation of three polar glycolipids - M5-DLO, MPD and GPD - from the cytoplasmic to the luminal leaflet of the ER (Sanyal, S. and Menon, A.K. 2009a, Schenk, B., Fernandez, F., et al. 2001). Biochemical assays provide compelling evidence that these transport events do not require metabolic energy (they are ATP-independent) and that they are mediated by ER membrane proteins with exquisite substrate specificity (Rush, J.S., van Leyen, K., et al. 1998, Rush, J.S. and Waechter, C.J. 1995, Sanyal, S., Frank, C.G., et al. 2008, Sanyal, S. and Menon, A.K. 2009b, Sanyal, S. and Menon, A.K. 2010). The molecular identity of these proteins (scramblases) is not known (Pomorski, T.G. and Menon, A.K. 2016, Sanyal, S. and Menon, A.K. 2009a).

The M5-DLO scramblase is critically important as *N*-glycosylation cannot occur without it. The ER membrane protein Rft1 was proposed as the M5-DLO scramblase almost two decades ago (Helenius, J., Ng, D.T., et al. 2002, Ng, D.T., Spear, E.D., et al. 2000), but subsequent work showed that whereas Rft1 is clearly important for *N*-glycosylation, it appears to have no direct role in translocating M5-DLO across the ER membrane (Frank, C.G., Sanyal, S., et al. 2008, Gottier, P., Gonzalez-Salgado, A., et al. 2017, Jelk, J., Gao, N., et al. 2013, Rush, J.S., Gao, N., et al. 2009, Sanyal, S., Frank, C.G., et al. 2008).

We developed methods to assay M5-DLO scramblase activity in large unilamellar vesicles reconstituted with a mixture of ER membrane proteins derived from yeast or rat liver. We used radiolabeled M5-DLO as the reporter lipid, and the α-mannosyl-binding lectin Concanavalin A as a topological probe (Frank, C.G., Sanyal, S., et al. 2008, Sanyal, S., Frank, C.G., et al. 2008, Sanyal, S. and Menon, A.K. 2009b, Snider, M.D. and Rogers, O.C. 1984, Vidugiriene, J. and Menon, A.K. 1994). The assay is described in detail in Fig. 1B. Using this assay, we showed that the M5-DLO scramblase activity is ATP-independent, and highly structure specific (Sanyal, S. and Menon, A.K. 2009a). Thus, higher order structures such as M7-DLO and M9-DLO are not scrambled efficiently (Sanyal, S., Frank, C.G., et al. 2008), nor is a structural isomer of M5-DLO in which the mannose residues correspond to the arrangement seen in ‘processed glycans’ (Sanyal, S. and Menon, A.K. 2009b). We were also able to resolve the M5-DLO scramblase from other scramblases responsible for transporting phospholipids and MPD (Sanyal, S., Frank, C.G., et al. 2008, Sanyal, S. and Menon, A.K. 2010). Despite these advances, a straightforward purification of the scramblase proved elusive indicating that a different approach may be useful.

## Results

### Identifying M5-DLO scramblase candidates by activity correlation profiling

Hypothesizing that M5-DLO scramblase activity is due to a single protein or protein complex, present as a homogeneous entity in the ER, we developed a mass spectrometry-based activity correlation profiling approach (McAllister, F.E. and Gygi, S.P. 2013, Sakurai, H., Kubota, K., et al. 2013) to identify scramblase candidates from a crude mixture of detergent-solubilized ER membrane proteins (Fig. 2). The underlying concept involves resolving the proteins in the crude mixture into a number of fractions using any separation technique of choice, measuring the scramblase activity of each fraction to generate an activity profile and, in parallel, using mass spectrometry to determine the relative abundance of individual proteins across the fractions (Fig. 2A). Proteins whose profiles best match the activity profile (Fig. 2B) are filtered through a data curation process to identify scramblase candidates.

**Figure 2.**
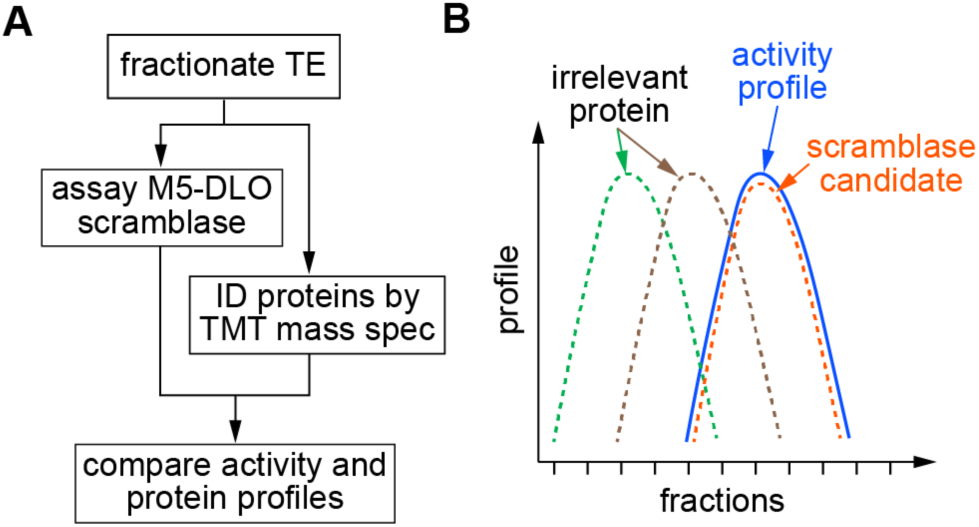
Principle of the activity correlation profiling approach. **A**.A Triton X-100 extract (TE) of salt-washed yeast membranes, i.e. an extract that is enriched in ER membrane proteins, is fractionated by velocity gradient sedimentation. Half of each fraction is reconstituted into large unilamellar vesicles and assayed for M5-DLO scramblase activity as outlined in Figure 1B to generate a profile of scramblase activity across the gradient. The remaining half of each fraction is subjected to mass spectrometric analysis using the tandem mass tag (TMT) system. Thus, each fraction is digested with trypsin, and the resulting peptides are labeled with TMT multiplex reagents (a unique TMT mass tag is used for each fraction). The digests are pooled and analyzed by tandem mass spectrometry to quantify the relative amount of specific proteins across fractions, thereby generating protein profiles. The protein profiles are quantitatively compared with the activity profile by obtaining a Pearson correlation score. **B**. Schematic illustration of the readout from the activity correlation profiling experiment. The activity profile (solid blue line) can be compared to the profile of several different proteins (dashed lines) identified by tandem mass spectrometry. Only a protein whose profile is highly correlated with the activity profile is considered as a scramblase candidate. Other proteins that fractionate differently (brown and green dashed lines) are considered irrelevant to scramblase activity.

Total membranes from a yeast cell homogenate were washed with high salt to remove peripheral proteins, and then treated with ice-cold Triton X-100 to extract ER membrane proteins as described previously (Sanyal, S., Frank, C.G., et al. 2008). The resulting ‘Triton Extract’ (TE) was fractionated by velocity gradient sedimentation using a continuous glycerol gradient. Sedimentation standards were analyzed in a parallel gradient. Gradient fractions were collected manually from the top and, after measurement of refractive index, the fractions were passed over a desalting spin column to remove glycerol. A portion of each fraction from the middle section of the gradient (fractions 5-11, shown in preliminary tests to contain the majority of M5-DLO scramblase activity) was reconstituted into large unilamellar vesicles and assayed for scramblase activity using [^3^H]M5-DLO (Fig. S1), whereas the remainder was taken for mass spectrometry using 6-plex tandem mass tag (TMT) reagents (Thompson, A., Schafer, J., et al. 2003).

The M5-DLO scramblase activity assay has been described in detail previously (Frank, C.G., Sanyal, S., et al. 2008, Sanyal, S., Frank, C.G., et al. 2008, Sanyal, S. and Menon, A.K. 2009b). The assay (Fig. 1B) exploits the organic-solvent resistant interaction between M5-DLO and the lectin Concanavalin A (Con A) to capture M5-DLO molecules located on the external surface of large unilamellar vesicles, and thereby separate them from those located in the inner leaflet. In the absence of a reconstituted functional scramblase, M5-DLO molecules located in the inner leaflet of the vesicles are inaccessible to Con A, whereas if the vesicles contain a scramblase then these molecules are translocated from the inner leaflet to the external leaflet and captured by the lectin. A key point is that the rate of scrambling is greater than the rate at which M5-DLO is captured by Con A (Sanyal, S., Frank, C.G., et al. 2008, Sanyal, S. and Menon, A.K. 2009b), even if the assay is conducted on ice, which would be expected to reduce the scrambling rate. Thus, the assay reports end-point data, ranging from 50-100% M5-DLO captured depending on the proportion of vesicles in the sample that is reconstituted with a functional M5-DLO scramblase. The reconstitution of M5-DLO scramblase molecules into an ensemble of lipid vesicles is governed by Poisson statistics (for example, see references (Pandey, K., Ploier, B., et al. 2017, Ploier, B., Caro, L.N., et al. 2016, Verchere, A., Ou, W.L., et al. 2017)). To know the amount of M5-DLO scramblase in a mixture of proteins, e.g. in a fraction from the velocity gradient, we would therefore have to carry out reconstitutions with a range of protein concentrations and use a Poisson model to calculate scramblase abundance. As this is an impractical undertaking when assaying fractions from the velocity gradient, we decided instead to estimate scramblase abundance by reconstituting just enough protein so that the fraction with the highest amount of M5-DLO scramblase would report approximately 70% M5-DLO capture. We determined empirically that this could be accomplished by reconstituting half of each fraction.

The quality of separation achieved on the velocity gradient is shown in Fig. 3A. The activity profile has a peak in fractions 7-8, corresponding to a nominal sedimentation coefficient of 6-7S, whereas the bulk of the protein peaks earlier in the gradient, in fraction 6. Approximately 3000 proteins were identified in the mass spectrometric analysis (Table S1 and Table S2). To learn how the various proteins were resolved on the gradient we determined the fraction in which each protein had its maximum abundance. This allowed us to obtain the average molecular weight of proteins that had their maximum abundance in each fraction. The data show that this average molecular weight increases from ∼40 kDa (fraction 5) to ∼80 kDa (fraction 8), remaining approximately constant at ∼80 kDa over fractions 8-10 (Fig. S2). This result is consistent with the general expectation that protein monomers and small complexes will predominate in the initial fractions and higher order complexes will be found in fractions 8-10. In line with this conclusion, the multi-subunit OST complex and the glycosyl-phosphatidylinositol (GPI) transamidase complex sediment relatively rapidly in the gradient, with the majority of their constituent subunits being maximally detected in fraction 7 (Fig. S3).

**Figure 3:**
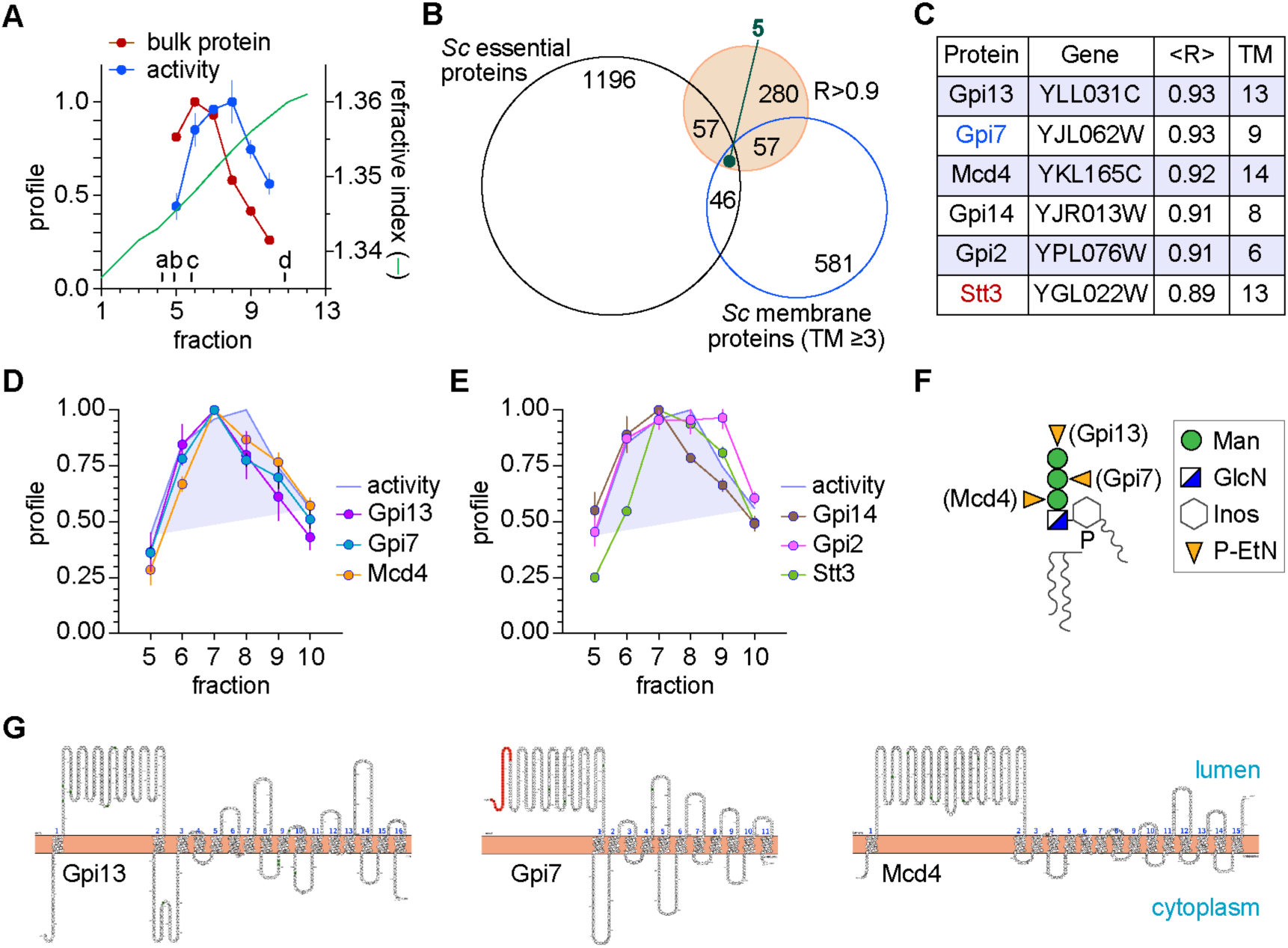
Identification of M5-DLO scramblase candidates by activity correlation profiling. **A**. Yeast TE was fractionated by velocity sedimentation on a 5-25% glycerol gradient and fractions were collected from the top. Sedimentation standards (a-d, corresponding to carbonic anhydrase (2.8S), ovalbumin (3.6S), bovine serum albumin (4.2S) and β-amylase (8.9S)) were resolved in a parallel gradient and their migration positions are indicated. The refractive index of the fractions is indicated by the green line. The profiles of bulk protein (carmine) and M5-DLO scramblase activity (blue, duplicate measurements) are indicated for fractions 5-10. **B**. Venn diagram describing the selection of M5-DLO scramblase candidates. The mass spectrometry analyses identified 280 proteins with a correlation coefficient R>0.9 (filled circle). Intersections between this group and the groups of all essential proteins in yeast (*Sc* essential proteins), and yeast membrane proteins with three or more predicted transmembrane spans (*Sc* membrane proteins (TM≥3, assessed by the TMHMM membrane protein topology prediction program)) yielded 5 proteins. One of these (Neo1) was discarded because it is not an ER resident. **C**. List of M5-DLO scramblase candidates. The four ER membrane proteins obtained at the intersection of the Venn diagram are listed. The non-essential protein Gpi7 was added because its profile is highly correlated with that of M5-DLO scramblase activity (see panel D) and because it belongs to the family of GPI phosphoethanolamine transferases that includes the identified candidates Gpi13 and Mcd4 (see panels F and G). Stt3 was added to the list despite R<0.9 because of its relevance to *N*-glycosylation as the catalytic subunit of OST (Figure 1A). **D**. Profile of candidates: profiles of Gpi13, Gpi7 and Mcd4 versus the activity profile (shaded and outlined in blue). **E**. Profile of candidates: profiles of Gpi14, Gpi2 and Stt3 versus the activity profile (shaded and outlined in blue). **F**. Role of Gpi13, Gpi7 and Mcd4 in P-EtN transfer reactions of GPI biosynthesis. The structure of the mature GPI anchor precursor in yeast and mammals is shown, with phospho-ethanolamine residues (P-EtN) decorating each of the mannose residues. Mcd4, Gpi7 and Gpi13 are the P-EtN transferases responsible for adding the first, second and third P-EtN residues, respectively. *T. brucei* lacks homologs of Mcd4 and Gpi7, and consequently has only the capping P-EtN residue transferred by Gpi13. **G**. Membrane topology of Gpi13, Gpi7 and Mcd4 assessed using Protter (Protter: interactive protein feature visualization and integration with experimental proteomic data. Omasits U, Ahrens CH, Müller S, Wollscheid B. Bioinformatics. 2014 Mar 15;30(6):884-6. doi: 10.1093/bioinformatics/btt607.) The salmon-coloured strip is the ER membrane bilayer. The stretch of Gpi7 coloured red is a predicted signal sequence. The lumenal and cytoplasmic sides of the membrane are indicated. The number of predicted transmembrane spans differs slightly from the number predicted by the TMHMM server that was used to calculate the Venn diagram group ‘*Sc* membrane proteins (TM≥3)’.

We next quantitatively compared the protein abundance profiles (obtained from each of two technical replicates (Table S1 and Table S2)) with the scramblase activity profile, identifying 280 proteins whose profile matched that of the activity profile with a correlation score of R ≥ 0.9 (Fig. S4). We subjected this list to several data curation steps. Hypothesizing that the *N*-glycosylation scramblase is an essential, ER-localized membrane protein, we discarded proteins that did not meet these criteria. Thus, 62 of the 280 proteins are known to be essential in yeast, and 5 correspond to ER-localized polytopic membrane proteins with 3 or more transmembrane spans as predicted by the TMHMM web server (Fig. 3B). These latter proteins are listed in Fig. 3C, and their profiles in comparison to the activity profile are shown in Figs. 3D, 3E. We included the highly correlated but non-essential protein Gpi7 because it belongs to the same GPI phospho-ethanolamine-transferase (P-EtN) family as Mcd4 and Gpi13 (Galperin, M.Y. and Jedrzejas, M.J. 2001, Orlean, P. and Menon, A.K. 2007) (Fig. 3F, Fig. S4) that were identified in our analysis; all three are polytopic membrane proteins with a large lumenal loop (Fig. 3G). Remarkably, the five candidates identified in our analyses are enzymes of the ER-localized GPI biosynthetic pathway (Orlean, P. and Menon, A.K. 2007), and three of these (Gpi13, Gpi14, Gpi2) are ubiquitously found in eukaryotes. We noticed that the profile of Stt3, the catalytic subunit of OST (Fig. 1A), matched closely with the activity profile (correlation score of 0.89, close to our cut-off value of 0.9). Because of its importance to *N*-glycosylation we included Stt3 in our list of candidates.

### Biochemical test of M5-DLO scramblase candidates

To determine whether the six candidates (Gpi13, Gpi7, Mcd4, Gpi14, Gpi2 and Stt3) play a role in M5-DLO scrambling, we determined whether the M5-DLO scramblase activity present in TE could be eliminated by quantitatively and specifically immunodepleting the candidate from the TE prior to reconstitution and assay. We previously used a similar approach to show that opsin accounts for all the phospholipid scramblase activity in detergent-solubilized retinal disc membranes (Menon, I., Huber, T., et al. 2011), and that Rft1 does not contribute to M5-DLO scramblase activity in TE (Frank, C.G., Sanyal, S., et al. 2008).

We obtained haploid yeast strains in which the M5-DLO scramblase candidate of interest is functionally expressed from its chromosomal locus with a C-terminal Tandem Affinity Purification (TAP) tag (Rigaut, G., Shevchenko, A., et al. 1999). The TAP-tagged protein is thus the only version of that protein being expressed in the cell. TE was prepared from strains expressing individual TAP-tagged M5-DLO scramblase candidates and either mock-treated, or treated with IgG resin to eliminate the TAP-tagged protein. Western blotting indicated that the TAP-tagged candidates were quantitatively removed by IgG-resin treatment, whereas an irrelevant ER membrane protein - Dpm1 - was still present in the TE (Fig. 4). We reconstituted equivalent amounts of mock-treated and IgG-treated TE into vesicles for M5-DLO scramblase activity assay. As our assay provides an end-point readout that reports on the proportion of vesicles that is equipped with a functional scramblase (see above), we expected to see a complete loss of activity for one of the candidates, thus pointing to its role in scrambling M5-DLO. Surprisingly, scramblase activity was not significantly affected by the removal of any of the candidates (Fig. 4). Thus, none of the candidates contributes significantly to the scramblase activity of TE. However, as described below, we were nevertheless interested in further exploring the possible function of the GPI P-EtN transferases in M5-DLO scrambling.

**Figure 4:**
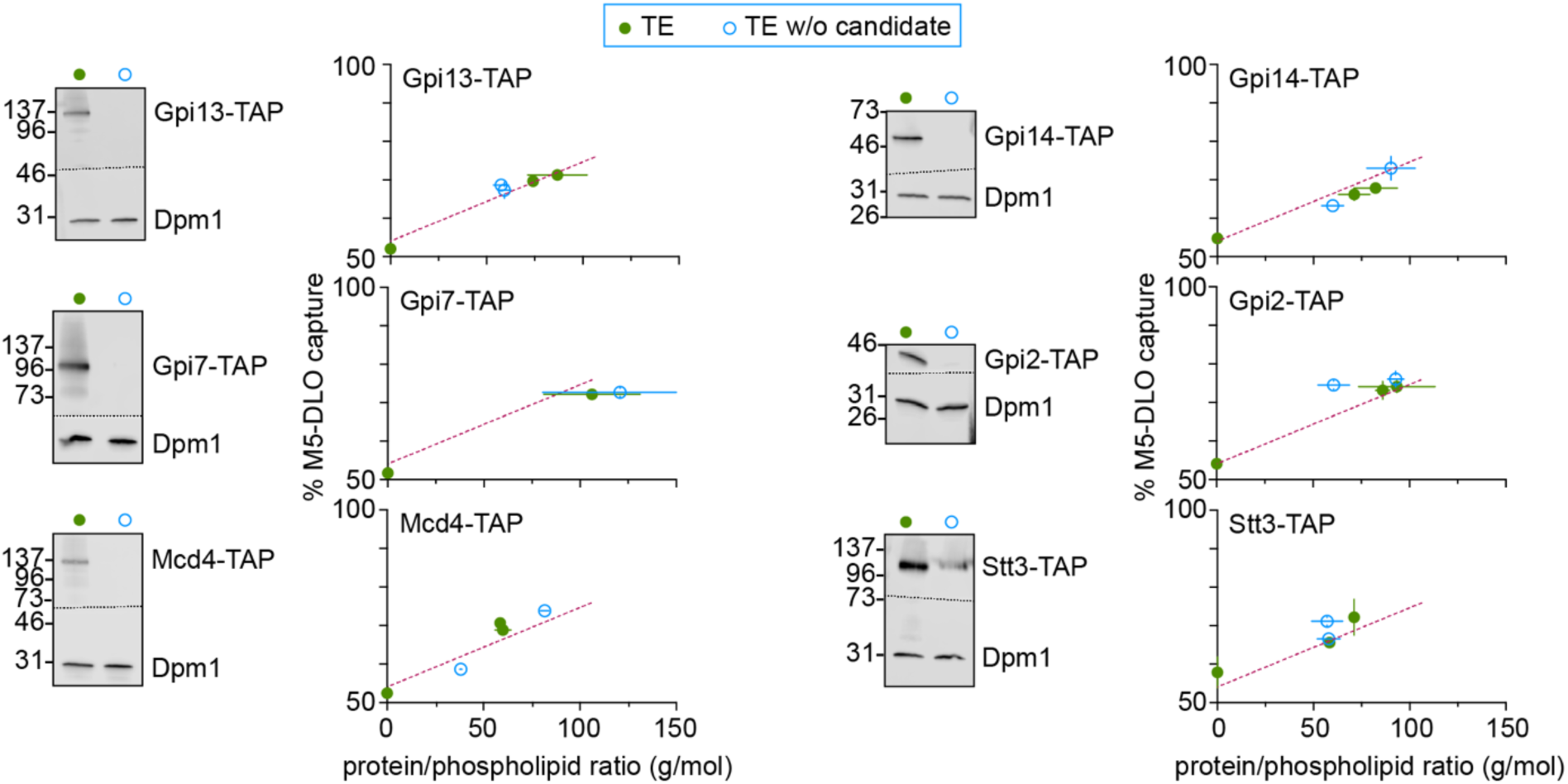
Evaluating M5-DLO scramblase candidates by immunodepletion. TE was prepared from yeast strains expressing a C-terminally TAP-tagged variant of the scramblase candidate in place of the endogenous protein. Tagging was done at the genomic locus. Six unique TE preparations were thus generated, corresponding to cells expressing Gpi13-TAP, Gpi7-TAP, Mcd4-TAP, Gpi14-TAP, Gpi2-TAP or Stt3-TAP. In every case the extract was mock-treated (green filled circles) or incubated with IgG-agarose beads to remove the TAP-tagged protein (blue open circles). Removal of the TAP-tagged protein from the extract was confirmed by SDS-PAGE/immunoblotting using anti-TAP antibodies, and specificity of immunodepletion and equivalence of sample loading was confirmed by immunoblotting to detect the ER resident protein Dpm1. Mock-treated and depleted extracts were reconstituted and assayed for M5-DLO scramblase activity. Reconstitutions were performed at different ratios of protein to phospholipid, in the approximate range 25 - 125 g/mol. The figure shows immunoblots paired with the corresponding activity assays for each candidate. Error bars are obtained from duplicate measurements of activity, as well as duplicate measurements of the protein and phospholipid content of the reconstituted vesicles. The red dashed line, based on all data from the mock-treated samples, is intended to guide the eye.

### Re-consideration of the GPI P-EtN transferase proteins as scramblase candidates

The three GPI P-EtN transferases have very similar membrane topology (Fig. 3G), carry out similar reactions (they each transfer P-EtN from phosphatidylethanolamine (PE) to mannose residues in the GPI core structure (Fig. 3F)), and are present at equivalent levels in the cell (Fig. S5). Previous reports indicate that they have functions in the cell distinct from their recognized roles in the GPI biosynthetic pathway. Thus, over-expression of any of the three proteins in yeast unexpectedly results in enhanced secretion of ATP, consistent with the possibility that each of these proteins is involved in the transport of ATP into the lumen of secretory compartments (Zhong, X., Malhotra, R., et al. 2003). Furthermore, Mcd4 was shown to have a non-canonical role in aminophospholipid metabolism (Storey, M.K., Wu, W.I., et al. 2001) and ER protein quality control (Ihara, S., Nakayama, S., et al. 2017), distinct from its role in GPI anchoring. Based on these indicators, we considered the possibility that the three GPI P-EtN transferases may each (redundantly) have M5-DLO scramblase activity. As the proteins are expressed at similar levels (Fig. S5), elimination of any one of them would result in a lowering of our scramblase assay read-out by one-third, whereas elimination of all three would result in total removal of scramblase activity from the TE. Although we did not see any significant difference in the assay readout when testing TE samples from which individual P-EtN transferases had been eliminated (Fig. 4), this could be a result of the potentially poor sensitivity of the assay to changes that are less than 2-fold. As described next, rather than continue with the reconstitution approach to explore a possible role of P-EtN transferases in M5-DLO scrambling, we took a genetic approach.

### *In vivo* test of Gpi13 as an M5-DLO scramblase

At the level of a single eukaryotic cell, *N*-glycosylation is essential for cell viability whereas GPI anchoring is generally not essential. Thus, mammalian cells in culture are non-viable when *N*-glycosylation is disrupted (Kelleher, D.J. and Gilmore, R. 2006), but their growth is unaffected by a deficiency in GPI anchoring (Hyman, R. 1988). The same holds true for the early diverging eukaryote *Trypanosoma brucei* (Guther, M.L., Lee, S., et al. 2006, Hong, Y., Nagamune, K., et al. 2006, Lillico, S., Field, M.C., et al. 2003, Nagamune, K., Nozaki, T., et al. 2000). Interestingly, GPI anchors in *T. brucei* have only a single P-EtN, linked to the third mannose residue, that provides the means to attach the anchor to proteins. Thus *T. brucei* has an ortholog of Gpi13 (TbGPI13), but lacks orthologs of Mcd4 and Gpi7. To test whether Gpi13 is essential for M5-DLO scrambling, we decided to generate TbGpi13 null mutants of insect stage (procyclic) *T. brucei*. We predicted that if TbGPI13 is solely involved in GPI anchoring, then *T. brucei* GPI13Δ cells would be viable. However, the *T. brucei* GPI13Δ cells would be inviable if TbGPI13 played an essential role in scrambling M5-DLO for protein *N*-glycosylation.

We sequentially disrupted the two alleles of TbGPI13 in procyclic-stage *T. brucei* using hygromycin and geneticin drug resistance cassettes (Fig. 5A) and recovered viable GPI13Δ cells that grew well, albeit approximately 2-fold more slowly than wild-type cells (Fig. 5B). Although our ability to generate viable GPI13Δ cells immediately indicates that Gpi13 does not play an essential role in *N*-glycosylation, we nevertheless characterized the cells to verify the expected disruption of GPI anchoring and to investigate the *N*-glycosylation status of specific glycoproteins. We performed metabolic radiolabeling with [^3^H]ethanolamine, and analyzed both polar lipids and proteins. Whereas both PE and the P-EtN-containing GPI anchor precursor PP1 (Field, M.C., Menon, A.K., et al. 1991) were labeled with [^3^H]ethanolamine in wild-type (WT) cells, PP1 was not labeled in GPI13Δ cells (Fig. 5C). Consistent with this observation, SDS-PAGE/fluorography of protein extracts showed radiolabeled GPI-anchored GPEET procyclin (arrow head) in WT but not GPI13Δ cells (Fig. 5D). Of note, radiolabeling of eukaryotic elongation factor 1A, a protein that is modified by ethanolamine phosphoglycerol (Signorell, A., Jelk, J., et al. 2008), was similar in both WT and GPI13Δ cells (Fig. 5D).

**Figure 5:**
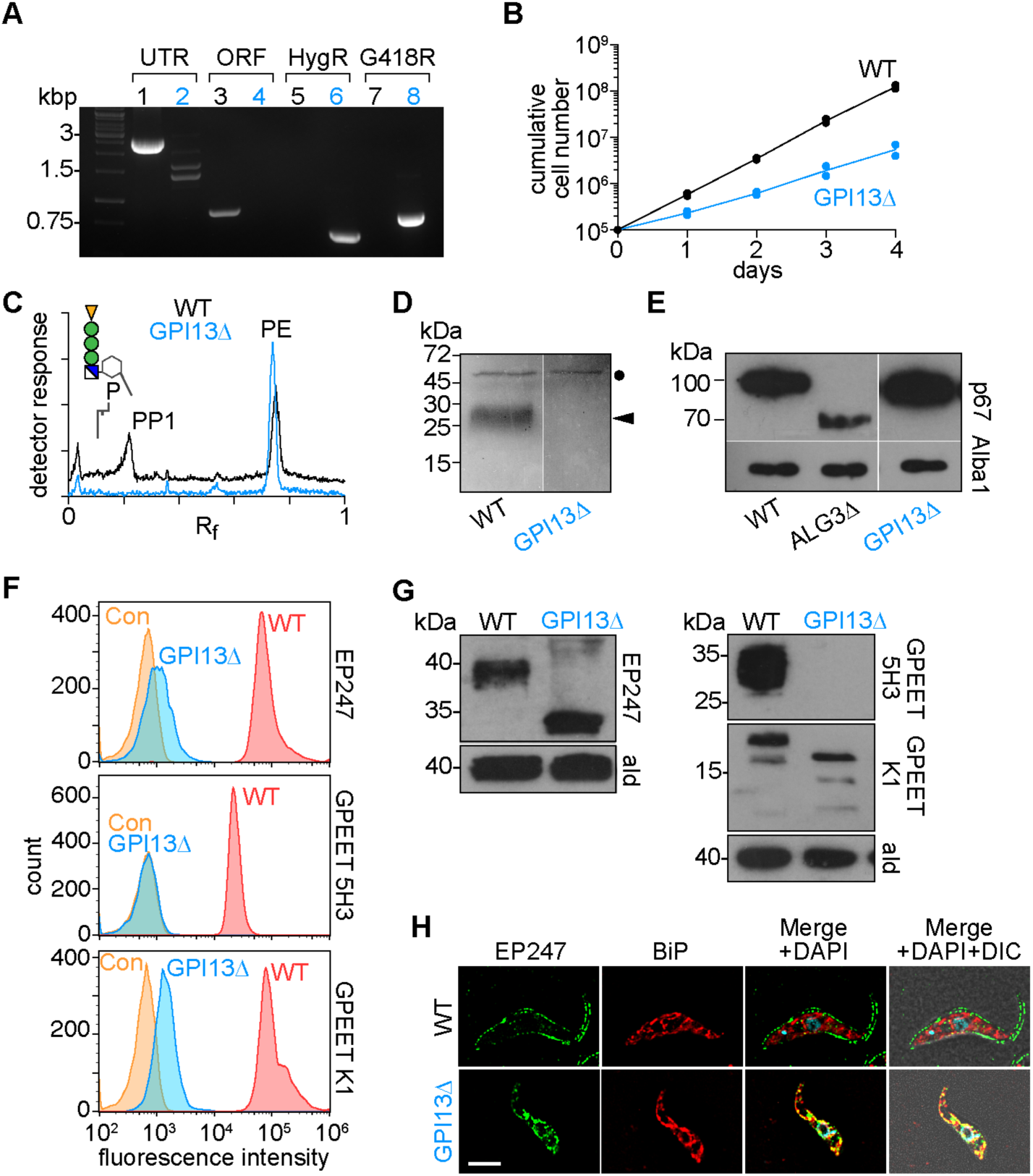
Evaluating Gpi13 as the M5-DLO scramblase in procyclic form *Trypanosoma brucei*. **A**. PCR confirmation of *TbGPI13* gene replacement using primers specific to the *TbGPI13* UTR and ORF, and HygR/G418R gene replacement cassettes. Lanes 1, 3, 5 and 7 correspond to wild-type cells, whereas lanes 2, 4, 6 and 8 correspond to TbGPI13 knockout cells. **B**. Growth of TbGPI13 knockout cells (GPI13Δ) compared with wild-type (WT) cells. Data points represent mean values from 2 independent experiments. **C**. Thin layer chromatographic analysis of polar lipid extracts from [^3^H]-ethanolamine-labeled WT (top trace) and GPI13Δ (bottom trace) cells showing loss of GPI synthesis precursor PP1 in GPI13Δ cells. The chromatograms were visualized using a radioactivity scanner. The WT trace is displaced upwards for clarity. PP1 and PE refer to the major GPI precursor in procyclic trypanosomes (structure shown in schematic, corresponding to EtN-P-Man_3_GlcN-(2-acyl-Inositol)-P-monoacylglycerol) and PE, respectively. The majority of PE is removed by extraction with a lower polarity solvent, before extracting PP1 in a more polar solvent. **D**. SDS-PAGE/fluorography of protein extracts from [^3^H]-ethanolamine-labeled WT and GPI13Δ cells showing loss of GPI-anchored GPEET procyclin (indicated by arrow head) in GPI13Δ cells. The band at ∼50 kDa (indicated by filled circle) represents eukaryotic elongation factor 1A, a protein that is modified by ethanolamine phosphoglycerol and therefore metabolically labeled with [^3^H]-ethanolamine (Signorell, A., Jelk, J., Rauch, M., Bütikofer, P. (2008) Phosphatidylethanolamine is the precursor of the ethanolamine phosphoglycerol moiety bound to eukaryotic elongation factor 1A. J. Biol. Chem. 283: 20320-20329) **E**. Analysis of p67 *N*-glycosylation by anti-p67 immunoblot of whole cell protein samples from WT and GPI13Δ cells. A sample from a known *N*-glycosylation defective cell line (ALG3Δ) was used as a positive control. Alba1 was probed using anti-Alba1 antibody and used as a loading control. **F**. Flow cytometry analysis of surface expression of GPI-anchored EP and GPEET procyclins in WT and GPI13Δ cells. The middle and right panels correspond to flow cytometry analysis of GPEET using 5H3 antibody (middle) and K1 antiserum (right). Control (Con) samples were generated by omitting the primary antibody. **G**. Anti-EP and anti-GPEET (K1) procyclin immunoblots of whole cell samples from WT and GPI13Δ cells. Aldolase was probed using anti-aldolase antibody and used as a loading control. **H**. WT and GPI13Δ cells were fixed and stained with antibodies against EP and BiP (as a marker for the endoplasmic reticulum) in combination with fluorophore-conjugated secondary antibodies and analyzed by fluorescence microscopy. DNA was stained with 4’,6-diamidino-2-phenylindole (DAPI) in the merged panels. DIC, differential interference contrast. Scale bar = 5 µm.

We tested the intactness of the *N*-glycosylation pathway in GPI13Δ cells by assessing the glycosylation status of the lysosomal *N*-glycoprotein p67 which has 14 *N*-glycosylation sites. An anti-p67 immunoblot of whole cell protein samples from WT and GPI13Δ cells revealed no change in the molecular mass of p67, running at approximately 100 kDa, consistent with normal *N*-glycosylation (Alexander, D.L., Schwartz, K.J., et al. 2002). In contrast, the mass of the glycoprotein was lower in a control *N*-glycosylation defective cell line (ALG3Δ) that cannot mannosylate M5-DLO *en route* to synthesis of the optimal DLO for the OST reaction (Manthri, S., Guther, M.L., et al. 2008) (Fig. 5E).

Finally, we examined the status of non-GPI-anchored procyclins in the GPI13Δ cells. We found that the main consequence of the lack of GPI anchoring was that the proteins remain sequestered inside the cell, failing to exit the ER as evinced by flow cytometry and fluorescence microscopy (Fig. 5F-H).

We conclude that GPI anchoring is disrupted as expected in GPI13Δ cells, whereas *N*-glycosylation is unperturbed. The inability to synthesize GPI-anchored surface coat proteins may explain the somewhat slower growth rate of the cells (Hong, Y., Nagamune, K., et al. 2006, Nagamune, K., Nozaki, T., et al. 2000)(Fig. 5B). Taken together with the biochemical data presented in Fig. 4, these results show that Gpi13 is not essential for M5-DLO scrambling as its deficiency does not impact *N*-glycosylation in a living cell.

## Discussion

We hypothesized that M5-DLO scramblase activity is essential in eukaryotes, and that if it is due to a single protein, then that protein would necessarily be essential for cell viability. Using activity correlation profiling (Fig. 2) in conjunction with a data filtering strategy (summarized in Fig. 3B) to implement this idea, we generated a list of 6 M5-DLO scramblase candidates (Fig. 3C). All six proteins are ER residents with multiple transmembrane spans, and they have well-established functions in GPI anchoring and protein *N*-glycosylation. However, our biochemical data indicate that none of these proteins contributes significantly to M5-DLO scramblase activity. Thus, quantitative removal (using genomic TAP-tagging and immunodepletion) of any one of the candidates from a crude mixture of ER membrane proteins (TE) prior to membrane reconstitution did not decrease the potency of the TE to populate an ensemble of large unilamellar vesicles with M5-DLO scramblase (Fig. 4).

Despite the negative outcome of our biochemical tests, we were especially interested in the family of three GPI P-EtN transferases - Gpi13, Gpi7 and Mcd4 - that we had identified as scramblase candidates. These proteins catalyze essentially identical reactions (Fig. 3F), and have a similar membrane topology comprising multiple transmembrane spans (Fig. 3G). Of considerable interest, all three were shown to participate in a function distinct from their role in GPI anchoring, i.e. ATP secretion (Zhong, X., Malhotra, R., et al. 2003). In addition, Mcd4 was shown to have a non-canonical role in aminophospholipid metabolism and ER protein quality control (Ihara, S., Nakayama, S., et al. 2017, Storey, M.K., Wu, W.I., et al. 2001). These intriguing moonlighting functions attributed to the GPI P-EtN transferases raised the possibility that the three proteins could potentially each, independently, contribute to M5-DLO scramblase activity. Our inability to detect these activities biochemically could be due to an inherent lack of sensitivity to the assay as performed. Thus, in order to identify the potential contributions of each of these equally expressed proteins (Fig. S5), we would have to (i) perform the assay over a range of protein-phospholipid ratios, and (ii) use a Poisson model to detect the change in abundance of M5-DLO scramblase after biochemical elimination of any one of the three proteins. Alternatively, we would have to purify each of the proteins and test for scramblase activity after reconstitution. Rather than carry out this latter sufficiency test which brings with it a unique set of issues, e.g. the protein may not be amenable to purification in functional form, we opted for a genetic approach. Knowing that *T. brucei* expresses only the Gpi13 ortholog of this 3-member family, we chose to knockout TbGPI13 and ask if we could recover viable cells. Our prediction was that if Gpi13 played an essential role in *N*-glycosylation, i.e. it is the sole M5-DLO scramblase in *T. brucei*, then we would not be able to generate GPI13Δ trypanosomes (as noted above, the GPI anchoring function of Gpi13 is not essential in *T. brucei* cells grown in culture)(Guther, M.L., Lee, S., et al. 2006, Hong, Y., Nagamune, K., et al. 2006, Lillico, S., Field, M.C., et al. 2003, Nagamune, K., Nozaki, T., et al. 2000). To our disappointment, we were able to generate viable GPI13Δ cells that had the expected deficiency in GPI anchoring (Fig. 5C, D), but no evident disruption of protein *N*-glycosylation (Fig. 5E). Thus, Gpi13 does not play an essential role in *N*-glycosylation (similar gene knockouts were also done for TbGPI2 and TbGPI14, resulting in viable cells (AJ, AC, PB, unpublished results) indicating that these proteins do not play an essential role in *N*-glycosylation).

It remains to discuss why the activity correlation profiling approach did not yield the identity of the M5-DLO scramblase. There are several possibilities.

i. We detected the vast majority (255) out of 286 known yeast ER membrane proteins (M. Schuldiner, unpublished data) (the 31 proteins that we did not detect are listed in Table S3), but the M5-DLO scramblase may have eluded detection. This could be because it might have only a few lysine and/or arginine residues, and that the resulting long tryptic peptides are not optimal for detection in the mass spectrometer. The relative resistance of M5-DLO scramblase activity to trypsin treatment (Sanyal, S., Frank, C.G., et al. 2008) is consistent with this possibility.
ii. We filtered the mass spectrometric data by imposing the criterion that M5-DLO scramblase candidates must have ≥ 3 TM spans. Known phospholipid scramblases of the GPCR and TMEM16 protein families have at least 7 TM spans (Goren, M.A., Morizumi, T., et al. 2014, Malvezzi, M., Chalat, M., et al. 2013, Menon, I., Huber, T., et al. 2011), consistent with their function as transporters. Indeed, if the M5-DLO scramblase would have fewer than 3 TM spans we would predict that it must homo- or hetero-oligomerize in order to form a functional transport entity.
iii. We hypothesized that M5-DLO scramblase activity is essential, and extrapolated this to imply that the activity is due to a single protein and that this protein would therefore also be essential. We considered the possibility that several proteins might individually possessing M5-DLO scramblase activity. In this case, the individual proteins may not be essential. Accordingly, we compiled a list of non-essential proteins whose profiles correlate well with the activity profile (Table S4). This list includes 18 ER-localized non-essential proteins with ≥ 3 predicted TM spans, plus 11 other proteins (also with ≥ 3 TM spans) whose subcellular location is not known. It would be interesting in the future to explore the possibility that these proteins may contribute to M5-DLO scramblase activity.
iv. Perhaps the most significant reason for the failure of the activity profiling approach is that the M5-DLO scramblase may co-exist in two or more forms, a complication that could account for the difficulty (noted in the Introduction) in purifying it. For example, the scramblase (Protein X) co-exists as a monomer with scramblase activity, and as a heterodimer (complexed with Protein Y) that is inactive. The protein profile of Protein X would thus track the rising portion of the activity profile (corresponding to monomeric Protein X) and then continue more broadly (corresponding to a complex of Protein X and Protein Y) before returning to baseline. The resulting broader profile of Protein X would result in it being discarded in our data curation as it would correlate poorly with the activity profile. Examples of such profiles (Fig. S6) reveal potentially interesting proteins such as the GlcNAc phosphotransferase Alg7 (R=0.65 ± 0)(mean ± SD) and the mannosyltransferase Alg2 (R= 0.66 ± 0.05) (mean ± SD). Both proteins recognize dolichyl diphosphate and could potentially play a role in scrambling M5-DLO. The profiles of both proteins track the rising portion of the activity profile but then broaden out even as the activity decreases. Additional scenarios are possible, for example that scramblase activity is due to a complex of two proteins that are individually inactive – in this situation the protein profiles would diverge from the rising portion of the activity profile, and only track with the activity as it decreases in higher fractions.

In conclusion, we describe an innovative strategy to identify the M5-DLO scramblase. Although the approach was ultimately unsuccessful, we report an extensive data set that provides an opportunity to extend these studies in the future. Our results raise the important possibility that the M5-DLO scramblase may be a protein, or protein complex, whose activity is regulated at the level of quaternary structure. This key point will be useful not only in future attempts to identify this enigmatic protein, but also in approaches aimed at the molecular identification of ER proteins involved in the transbilayer movement of MPD and GPD.

## Materials and methods

### Materials

Analytical-grade reagents were purchased from Sigma-Aldrich unless stated otherwise. Antibiotics were from Sigma-Aldrich, Invivogen or Invitrogen. Other reagents/materials were obtained as follows: D-2-[^3^H]-mannose (20-30 Ci/mmol) (American Radiolabeled Chemicals Inc.), Micro BCA protein assay kit (Thermo Fisher Scientific), Triton X-100 (Roche), IgG-Agarose resin (GE Healthcare), anti-TAP antibody (Genescript), anti-Dpm1 antibody (Abcam), SM-2 Biobeads and P6-Biogel resin (Biorad) and egg phosphatidylcholine (Avanti Polar Lipids). Yeast strains used in this study are listed in Table S5.

### Buffers

Buffer A: 10 mM HEPES-NaOH pH 7.4, 100 mM NaCl, 1% (w/v) Triton X-100; Buffer B: 10 mM HEPES-NaOH pH 7.4, 100 mM NaCl; Buffer C: Buffer B supplemented with 3 mM MgCl_2_, 1 mM CaCl_2_, 1 mM MnCl_2_.

### Preparation of [^3^H]M5-DLO

#### Protocol 1

Procyclic form *T. brucei* lacking TbRft1 (TbRft1^-/-^) were generated from *T. brucei* strain Lister 427 and maintained in culture as previously reported (Gottier, P., Gonzalez-Salgado, A., et al. 2017). 10^9^ cells were washed with glucose-free SDM-80, resuspended at a density of 1 × 10^8^ cells/ml in the same medium, and incubated for 90 min at 27°C. D-[2-^3^H]-mannose (200 μCi) was added and the cells were incubated for a further 2 hr at 27°C. The cells were then collected by centrifugation and lipids were sequentially extracted from the washed cell pellet with 3 x 5 ml chloroform/methanol (3:2, v/v), 3 x 5 ml chloroform/methanol/water (3:48:47, v/v/v), containing 1 M MgCl_2_, and 1 x 5 ml chloroform/methanol/water (3:48:47, v/v/v) without MgCl_2_. Finally, [^3^H]M5-DLO was extracted with 2 x 5 ml chloroform/methanol/water (10:10:3, v/v/v). Extracts were pooled, dried under a stream of nitrogen, and partitioned between water and n-butanol. [^3^H]M5-DLO was recovered in the n-butanol phase.

#### Protocol 2

Yeast cells (*alg3*Δ) were metabolically labeled with D-[2-^3^H]mannose in low-glucose medium as previously described (Sanyal, S., Frank, C.G., et al. 2008), and subjected to differential solvent extraction as for Protocol 1 to purify [^3^H]M5-DLO.

#### Radiochemical purity of [^3^H]M5-DLO preparations

The radiochemical purity of [^3^H]M5-DLO preparations was evaluated by thin layer chromatography (TLC) of the intact lipid, and by HPLC analysis of the oligosaccharide portion. TLC was done on glass-backed Silica Gel 60 plates (Merck) with chloroform/methanol/water (10/10/3, v/v/v) as the solvent system. The chromatograms were visualized using a Raytest Rita* radioactivity TLC analyser (Berthold Technologies). HPLC analysis was carried out by Bobby Ng and Hudson Freeze (Sanford-Burnham-Prebys Medical Discovery Institute, La Jolla, CA) as described (Kim, S., Westphal, V., et al. 2000). Briefly, the [^3^H]M5-DLO preparation was dried and treated with a 1:1 (v/v) mixture of isopropanol and 0.2 N HCL for 30 min at 100°C. The released oligosaccharide was analyzed on a Dionex Ultimate 2000 HPLC system using an Agilent Microsorb NH2-column. Radioactivity was detected using an in-line Lablogic Radiodetector and the analysis was calibrated using a co-injected 2-aminobenzamide-labeled dextran ladder (ProZyme/Agilent) detected on a Dionex fluorescence detector.

### Triton extract (TE) enriched in yeast ER membrane proteins

TE was prepared from yeast cells as described previously (Chalat, M., Menon, I., et al. 2012), except that an additional step was included to salt-wash the membranes prior to detergent solubilization. Briefly, 600 OD_600_ units of cells (BY4741, or BY4741-derived strain expressing a TAP tagged version of the protein of interest) were harvested, washed and homogenized using glass beads. After low-speed centrifugation to clear unbroken cells, the homogenates were centrifuged at 200,000 x *g*_*av*_ for 30 min to pellet membranes. The membranes were salt-washed by resuspension in buffer containing 1 M sodium acetate, followed by incubation on ice for 30 min. After re-pelleting, the salt-washed membranes were resuspended in ice-cold Buffer B before being solubilized by gradually adding an equal volume of 2% (w/v) ice-cold Triton X-100 at a final concentration of 1 % (w/v). The sample was incubated on ice for 30 min before removing insoluble material by centrifugation at 200,000 x *g*_*av*_ for 1 hr to generate a clear supernatant (TE). We previously used SDS-PAGE immunoblotting to show that TE is enriched in ER membrane proteins such as Dpm1 and Sec61, and depleted of plasma membrane proteins (Pma1, Gas1), as well as proteins of the vacuole (Vph1) and Golgi/endosomes (Pep12) (Sanyal, S., Frank, C.G., et al. 2008).

### Fractionation of TE by velocity sedimentation on a glycerol gradient

1.5 mL TE was loaded onto a 10 mL continuous glycerol gradient (5% to 25% (w/v), prepared in Buffer A) and centrifuged for 30 h in an SW41 Ti rotor at 100,000 x *g*_*av*_. A mixture of sedimentation standards (anhydrase, ovalbumin, bovine serum albumin, β-amylase) was loaded onto a parallel gradient. Fractions were collected manually or using an Auto Densi-Flow apparatus (Labconco). The refractive index of each fraction was measured before removing glycerol by buffer exchange on a desalting spin column packed with Biogel-P6 resin equilibrated in Buffer A. Migration of the sedimentation markers was determined by SDS-PAGE/Coomassie-staining of the fractions.

### Immunodepletion of a candidate protein from TE

TE from yeast strains expressing a TAP-tagged protein of interest (Table S5) was divided into two aliquots. One aliquot was treated at 4°C for 2 h with IgG-agarose beads to remove specifically the TAP-tagged protein, whereas the other was mock-incubated. Removal of the protein from the extract was verified by SDS-PAGE immunoblotting, using an anti-TAP antibody (1:1,000 dilution) and anti-Dpm1 (1:1,000). The Dpm1 immunoblot served as a specificity control as well as a loading control.

### Quantitative mass spectrometry

Six samples (fractions 5 to 10 of the gradient) were analyzed by mass spectrometry as follows. Proteins in each sample were digested by trypsin after reduction and alkylation with dithiothreitol and iodoacetamide. The resulting peptides were labeled with 6-plex TMT reagents. The digests were combined and subjected to offline fractionation by high pH reversed phase chromatography to obtain 6 fractions. Each fraction was analyzed by LC-MS. Online chromatography was performed with a Thermo Easy nLC 1000 ultra-high-pressure HPLC system (Thermo Fisher Scientific) coupled online to an Orbitrap Fusion Lumos mass spectrometer with a NanoFlex source (Thermo Fisher Scientific). Analytical columns (∼ 25 cm long and 75 µm inner diameter) were packed in-house with ReproSil-Pur C18 AQ 3 µM reversed phase resin (Dr. Maisch GmbH, Ammerbuch-Entringen, Germany). A peptide mixture was loaded onto the analytical column with buffer A (0.1% formic acid) and separated with a linear gradient of 3% to 32% buffer B (100% ACN and 0.1% formic acid) at a flow rate of 300 nL/min over 240 min. MS data were acquired using a data-dependent method, dynamically choosing the most abundant not-yet-sequenced precursor ions from the survey scans. Peptide fragmentation was performed via higher energy collisional dissociation. The MS data were processed by the MaxQuant software for protein identification and quantitation. The false discovery rate for protein identification was 1%. A Pearson correlation score (R) was obtained for each identified protein by comparing the measured profile profile and the activity profile.

### Candidate selection

Proteins with R >0.9 were selected. Using Yeastmine (Balakrishnan, R., Park, J., et al. 2012) this list was intersected with the list of all membrane proteins (Vazquez, H.M., Vionnet, C., et al. 2016), as well as all essential proteins in *Saccharomyces cerevisiae* (list generated using Yeastmine). This final list was manually completed with the number of transmembrane domains and cellular localization for each protein using information available in the Saccharomyces Genome Database.

### Liposomes and proteoliposome preparation

Vesicles were reconstituted essentially as described (Frank, C.G., Sanyal, S., et al. 2008, Sanyal, S., Frank, C.G., et al. 2008). Briefly, for each reconstitution sample, [^3^H]M5-DLO (20,000 cpm) was mixed with 4.5 µmol egg PC, dried under a stream of N_2_ and resuspended in 540 µl Buffer B. The lipids were sonicated before being supplemented with Triton X-100 (from a 10 % stock solution in water, final concentration 1 % (w/v)). For proteoliposomes, TE or a fraction from the velocity gradient was added; for control (protein-free) liposomes, an equivalent amount of Buffer A was added instead. The final volume of the sample was 1 ml. Vesicle formation was induced by removing detergent with SM2 biobeads (previously washed with methanol and water) added in two stages: 100 mg of the beads were added and the sample was incubated with end-over-end mixing at room temperature; following this incubation, a further 200 mg of the beads were added and vesicles were incubated with end-over-end mixing overnight at 4°C. The resulting vesicles were pelleted by centrifugation (200,000 x *g*_*av*_, 1 hr, 4°C) and resuspended in Buffer C. Vesicle protein was estimated using the Kaplan Pederson method (Kaplan, R.S. and Pedersen, P.L. 1985) and phospholipid was measured as described previously (Rouser, G., Siakotos, A.N., et al. 1966).

### M5-DLO scramblase assay

The assay was performed as previously described (Frank, C.G., Sanyal, S., et al. 2008, Sanyal, S., Frank, C.G., et al. 2008, Sanyal, S. and Menon, A.K. 2009b). Briefly, 20 µL of vesicles (proteoliposomes or liposomes) were incubated in 40 µl of Buffer C (“buffer”), or Buffer C containing 3.6 mg/ml Concanavalin A (“ConA”) or Buffer E containing 3.6 mg/ml Concanavalin A and Triton X-100 1.5% (w/v) (“ConA-TX”) for 30 min on ice. Then 140 µl of blocking solution (BSA 20 mg/ml, Yeast Mannan 2 mg/ml) was added, followed immediately by 1.4 ml of chloroform/methanol (1:1, v/v). The sample was vortexed and centrifuged (20,000 x *g*_*av*_, 5 min, 4°C). The supernatant was placed in a 20-ml scintillation vial for drying and the pellet was solubilized in 1 % (w/v) SDS overnight under vigorous shacking. Radioactivity in the pellet (P) and the supernatant (S) was measured by liquid scintillation counting. For each sample the percentage of M5-DLO capture was calculated as follows: 100• ((P+S)_ConA_) - (P+S)_buffer_)/ ((P+S)_ConA-TX_ - (P+S)_buffer_).

### Generation of TbGPI13 and TbGPI14 knock out strains in *T. brucei* procyclic forms

Gene disruptions were accomplished in *T. brucei* Cas9-expressing SmOx p9 procyclic forms by replacing both alleles of the target gene with antibiotic resistance genes using the CRISPR/Cas9 technique as described previously (Beneke, T., Madden, R., et al. 2017). Geneticin and hygromycin replacement cassettes, flanked by 30 bp sequences homologous to the target gene UTRs, were PCR-amplified from pPOTv7 plasmids using gene-specific forward and reverse primers (Table S6). Short guide RNA (sgRNA) templates, consisting of a T7 RNA polymerase promoter sequence and a gene-specific sgRNA sequence to guide double-strand breaks to the UTRs of the target gene, were amplified by annealing gene-specific primers with the G00 sgRNA scaffold primer. All PCR products were purified using Promega Wizard SV PCR clean-up system before transfection. Gene-specific geneticin and hygromycin targeting fragments and 5’ and 3’ sgRNA templates were combined with 1 x TbBSF buffer (90 mM sodium phosphate, 5 mM potassium chloride, 0.15 mM calcium chloride, 50 mM HEPES, pH 7.3) in a total volume of 100 µl and transfected into 3 x 10^7^ SmOx p9 *T. brucei* cells by electroporation using the Amaxa Nucleofector 4D (Lonza), program FI-115. Clonal populations resistant to both hygromycin and geneticin were obtained by limiting dilution, and complete loss of the target gene was verified by diagnostic PCR reactions.

### SDS-PAGE and immunoblotting

*T. brucei* cells were harvested by centrifugation, washed twice with PBS and the pellet suspended directly in SDS sample buffer and heated at 55°C for 10 mins. Approximately 5×10^6^ cell equivalents per sample were separated by SDS-PAGE under reducing conditions on a 12% polyacrylamide gel and transferred to nitrocellulose membrane. For all blots, blocking was carried out in 5% milk in TBS, and primary and secondary antibodies diluted in 1% milk in TBS + 0.1% Tween-20 (TBST), with the addition of 0.01% SDS for the secondary antibodies. The following primary antibodies were used: anti-p67 (J. Bangs, 1:5000), anti-ALBA1 (I. Roditi, 1:2500), anti-EP247 (Cedarlane, 1:1500), anti-GPEET K1 (I. Roditi, 1:2500), anti-GPEET 5H3 (I. Roditi, 1:5000) and anti-aldolase (P. Michels, 1:180,000). HRP-conjugated secondary anti-rabbit and anti-mouse antibodies (DAKO) were used at 1:1000 and 1:5000 respectively, and chemiluminescent detection carried out using SuperSignal™ Pico West PLUS substrate (ThermoScientific).

### Flow cytometric analysis

Trypanosomes (∼5×10^6^ cells) were centrifuged and the pellet was directly incubated with one of the following primary antibodies, anti-EP247, GPEET K1 or anti-GPEET 5H3, in SDM-79 media for 30 min. After two washes with SDM-79 samples were incubated with anti-rabbit or anti-mouse AlexaFluor 488 (Invitrogen) secondary antibody (both 1:1000) in SDM-79 for 30 min. Samples were washed two more times and analyzed with a Novo-Cyte flow cytometer (ACEA Biosciences). Samples and buffers were kept on ice throughout, and all centrifugations and antibody incubations were performed at 4°C.

### Metabolic labeling and extraction of [^3^H]-ethanolamine-labeled GPI precursors and GPI-anchored proteins

Procedures were as previously described (Gottier, P., Gonzalez-Salgado, A., et al. 2017). Briefly, 5×10^8^ cells were cultured for 16 hours in the presence of 40 μCi of [^3^H]ethanolamine. The cells were harvested by centrifugation and washed twice with ice-cold TBS (10 mM Tris-HCl, 144 mM NaCl, pH 7.4). Bulk phospholipids were extracted from the pellet with 2x 10 ml of chloroform/methanol 2:1 (v/v). GPI precursors and free GPIs were extracted with 3x 10 ml of chloroform/methanol/water 10:10:3 (v/v/v), and GPI-anchored proteins were extracted with 2x 10 ml of 9% butanol in water (v/v). GPI precursors and free GPIs were further separated by partitioning the extract between butanol and water. GPI precursors were analyzed by TLC while free GPIs and GPI-anchored proteins were visualized by SDS-PAGE followed by fluorography.

## Acknowledgements

We thank Jennifer Jelk for technical assistance, Guoan Zhang and his colleagues at the Proteomics and Metabolomics Core Facility (Weill Cornell Medicine) for mass spectrometry, Maya Schuldiner (Weizmann Institute) for sharing a list of yeast ER membrane proteins, Bobby Ng and Hudson Freeze (Sanford-Burnham-Prebys Medical Discovery Institute, La Jolla, CA) for oligosaccharide analysis of [^3^H]M5-DLO preparations, Brenda Andrews (University of Toronto) and Markus Aebi (ETH Zürich) for yeast strains, and Jay Bangs (SUNY Buffalo), Isabel Roditi (Universität Bern) and Paul Michels (Research Unit for Tropical Diseases, International Institute of Cellular and Molecular Pathology) for antibodies.

## Supplementary Figures

**Figure S1.**
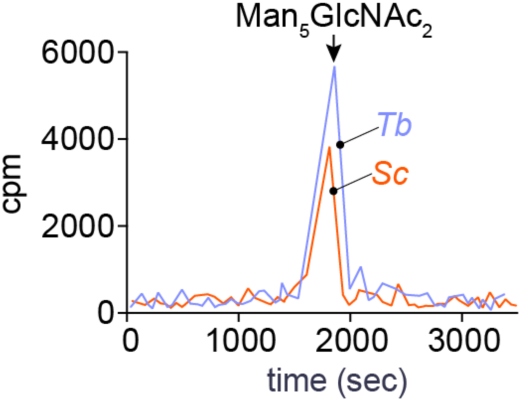
HPLC analysis of the oligosaccharide portion of [^3^H]M5-DLO. The pyrophosphate linkage in [^3^H]M5-DLO was hydrolyzed by mild acid treatment and the released [^3^H]oligosaccharide was analyzed by HPLC using an amino-column. Representative profiles (radioactivity (cpm), versus elution time) of samples of [^3^H]M5-DLO prepared via Protocol 1 (*Tb*) and Protocol 2 (*Sc*) are shown. The peak in each case co-elutes approximately with the glucose-9 oligomer in a dextran ladder that was co-injected with the samples. Data generated by Bobby Ng and Hudson Freeze.

**Figure S2.**
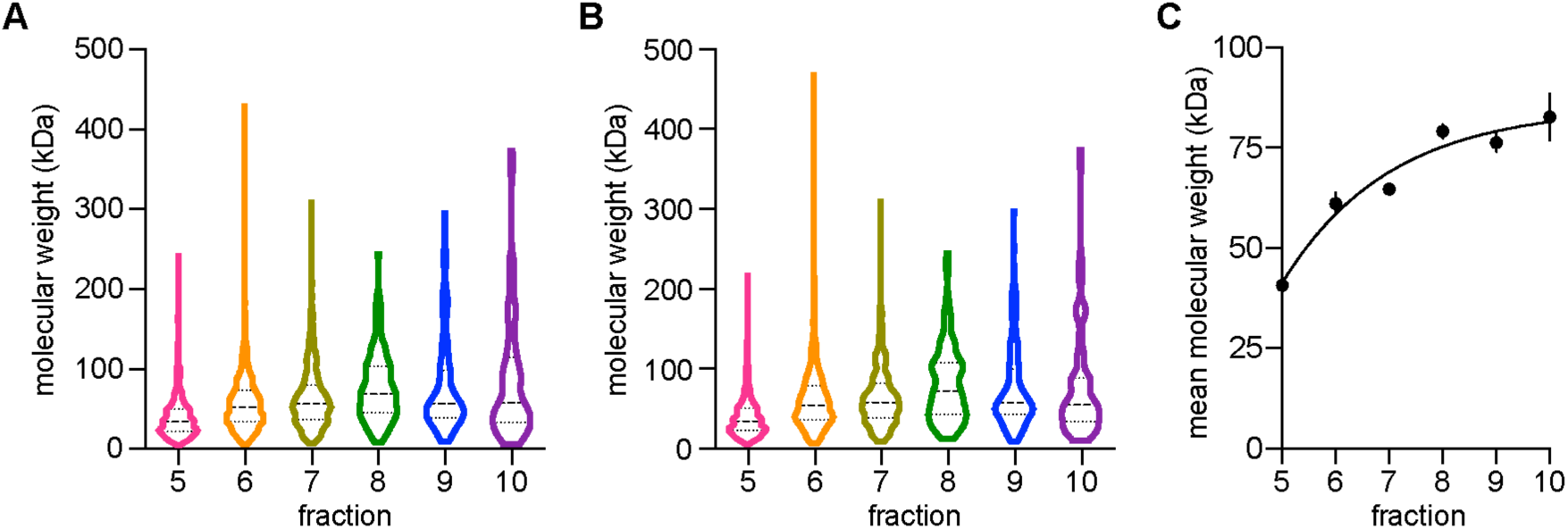
Distribution of proteins in the velocity gradient. The mass spectrometry data were used to determine the peak fraction for each identified protein, i.e. the fraction in which the protein is maximally abundant. The molecular weights of the proteins were obtained from the YeastMine webserver. **A, B**. Box and violin plots showing the molecular weights of proteins peaking in each fraction for the two technical replicates of the mass spectrometric analysis. The median and quartiles of the data are indicated. **C**. Mean molecular weights of proteins maximally abundant in each fraction. The points show the mean (± standard deviation) of data obtained from two technical replicates. The line through the points is intended to guide the eye.

**Figure S3.**
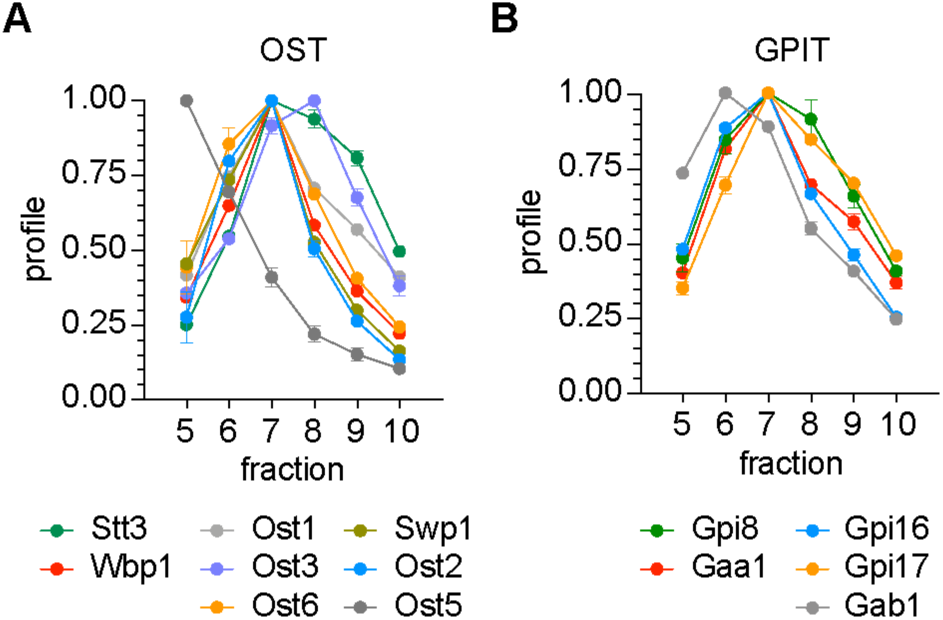
Profiles of OST and GPIT subunits in the velocity gradient. The distribution of subunits of the OST complex (panel A) and the GPI transamidase (GPIT) complex (panel B) in the velocity gradient were obtained from the mass spectrometry data. Yeast OST exists in two isoforms that contain either Ost3 or Ost6, in addition to seven other subunits: Stt3, Ost1, Wbp1, Swp1, Ost2, Ost4, Ost5. These complexes can be isolated from yeast membranes after solubilization in dodecylmaltoside detergent supplemented with cholesteryl hemisuccinate (Wild et al. *Science* **359**: 545-550 (2018)). In panel A, Triton X-100-solubilized OST complex remains largely intact, with the exception of the Ost5 subunit which sediments less rapidly than the rest of the subunits and appears to have dissociated from the complex.

**Figure S4:**
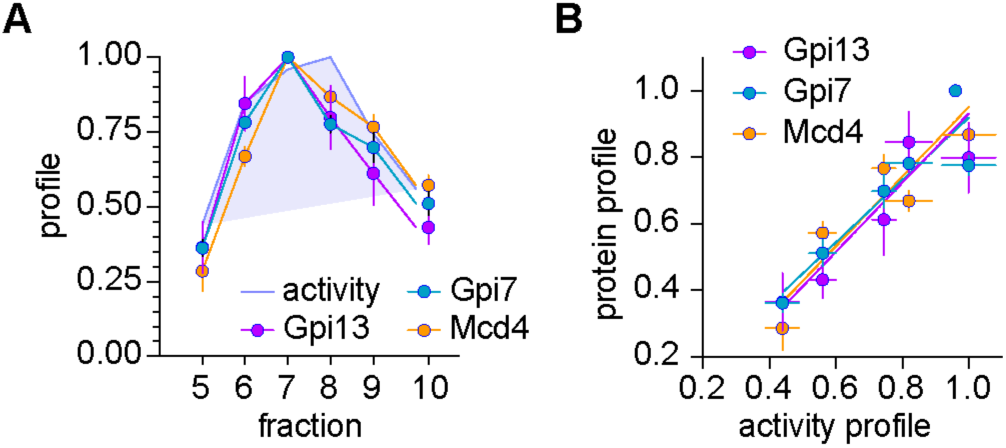
Correlation between the activity profile and the profiles of Gpi13, Gpi7 and Mcd4. **A**. Reproduced from Fig. 3D. **B**. Quantification of correlation between the activity profile and profiles of Gpi13, Gpi7 and Mcd4 (taken from panel A). Values defining the profile of a specific protein, i.e. relative protein abundance in a particular fraction, are plotted against the corresponding values of activity and analyzed by simple linear regression. Linear regression yielded R squared values of >0.81, corresponding to R values of >0.90. The plot shows intuitively that activity values are high when relative protein abundance values are high, and correspondingly that activity values are low when relative protein abundance values are low.

**Figure S5:**
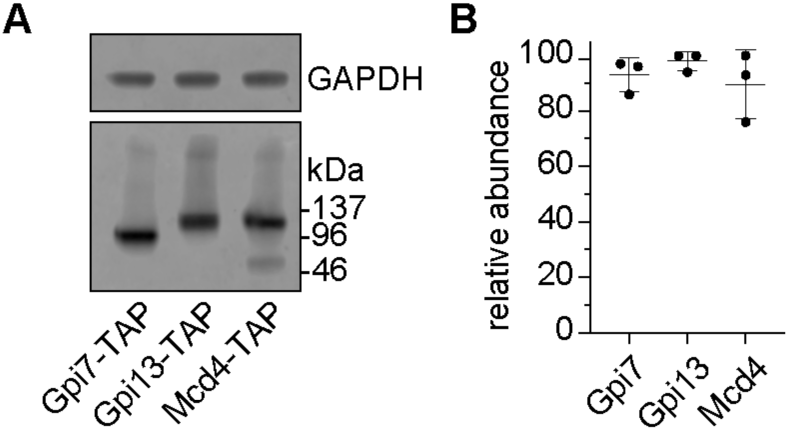
Gpi7, Gpi13 and Mcd4 are expressed at similar levels. **A**. 6 OD of cells expressing chromosomally TAP-tagged Gpi7, Gpi13 or Mcd4 were precipitated with TCA, pellets were washed with acetone, resuspended in loading dye and analyzed by SDS-PAGE/immunoblotting using anti-TAP antibody. Immunoblotting of GAPDH served as a loading control. **B**. The intensity of bands detected by anti-TAP and anti-GAPDH was quantified using Image Studio software. The ratio of intensities was taken as an indicator of abundance of the TAP-tagged protein. Data for three analyses are shown (mean ± standard deviation).

**Figure S6:**
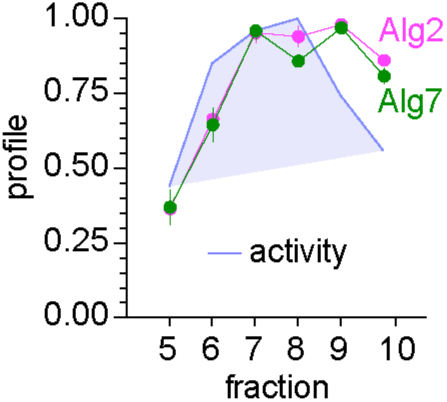
Profiles of Alg2 and Alg7. Profiles of Alg2 and Alg7 illustrate the possibility that the activity correlation profile method may be complicated by complex formation (see Discussion for details).

Table S1

**Table S2.** Table S2 Technical replicates of TMT mass spectrometric analysis of velocity gradient fractions (see Fig. 3A and ‘Materials and methods’ for details). The Tables are provided as .csv (comma separated value) files. A description of the column headings used in the Tables is given below.

| Column Name | Description |
| --- | --- |
| Protein IDs | Identifier(s) of protein(s) contained in the protein group. |
| Protein names | Name(s) of protein(s) contained within the group. |
| Gene names | Name(s) of the gene(s) associated to the protein(s) contained within the group. |
| FASTA headers | FASTA headers(s) of protein(s) contained within the group. FASTA file is a text-based format containing protein sequences, in which amino acids are represented using single-letter codes. |
| Peptides | The total number of peptide sequences associated with the protein group (i.e. for all the proteins in the group). |
| Sequence coverage [%] | Percentage of the sequence that is covered by the identified peptides of the best protein sequence contained in the group. |
| Q-value | This is an indicator of the confidence for protein identification, with range of 0-0.01 (1%). A smaller Q-value represents higher confidence of protein identification. |
| Columns H-M | Summed up intensity of all the identified peptide sequences |
| cor_score | The correlation score is the Pearson correlation between the measured profile and the standard profile (see Fig. S4) |

**Table S3.** Predicted ER membrane proteins that were not identified in the present study

| Gene | Protein | Function | TM spans |
| --- | --- | --- | --- |
| YLL055W | YCT1 | Yeast Cysteine Transporter | 10 |
| YLR004C | THI73 | THI regulon | 7 |
| YLR023C | IZH3 | Implicated in Zinc Homeostasis | 7 |
| YMR274C | RCE1 | Ras and a-factor Converting Enzyme | 7 |
| YHR140W |  |  | 6 |
| YER140W | EMP65 | ER Membrane Protein of 65 kDa | 5 |
| YKR053C | YSR3 | Yeast Sphingolipid Resistance | 5 |
| YFR042W | KEG1 | Kre6-binding ER protein responsible for Glucan synthesis | 4 |
| YLR246W | ERF2 | Effect on Ras Function | 4 |
| YNL008C | ASI3 | Amino acid Sensor-Independent | 4 |
| YNL194C |  |  | 4 |
| YLL014W | EMC6 | ER Membrane protein Complex | 3 |
| YIL089W |  |  | 2 |
| YFR041C | ERJ5 | Endoplasmic Reticulum located J-protein | 2 |
| YDR437W | GPI19 | Glycosyl Phosphatidylinositol anchor biosynthesis | 2 |
| YGR105W | VMA21 | Vacuolar Membrane Atpase | 2 |
| YLL052C | AQY2 | AQuaporin from Yeast | 2 |
| YNL046W |  |  | 2 |
| YNL146W |  |  | 2 |
| YOR044W | IRC23 | Increased Recombination Centers | 2 |
| YPL096C-A | ERI1 | ER-associated Ras Inhibitor | 2 |
| YAL028W | FRT2 | Functionally Related to TCP1 | 1 |
| YBL100C |  |  | 1 |
| YBR162W-A | YSY6 |  | 1 |
| YDL232W | OST4 | OligoSaccharylTransferase | 1 |
| YLR238W | FAR10 | Factor ARrest | 1 |
| YNL087W | TCB2 | Three Calcium and lipid Binding domains (TriCalBins) | 1 |
| YOR324C | FRT1 | Functionally Related to TCP1 | 1 |
| YPL200W | CSM4 | Chromosome Segregation in Meiosis | 1 |
| YER053C-A |  |  | 1 |
| YMR122W-A | NCW1 | Novel Cell Wall protein | 1 |

**Table S4.** Non-essential membrane proteins identified with R>0.9 and ≥3 TM spans

| ORF | Protein |  | MW (Da) | TM spans |
| --- | --- | --- | --- | --- |
| ER localized |  |  |  |  |
| YDR205W | MSC2 | Meiotic Sister-Chromatid recombination | 80578.2 | 13 |
| YBR132C | AGP2 | high-Affinity Glutamine Permease | 67261.4 | 12 |
| YJL212C | OPT1 | OligoPeptide Transporter | 91630.2 | 12 |
| YOR067C | ALG8 | Asparagine-Linked Glycosylation | 67409.4 | 12 |
| YGR227W | DIE2 | Derepression of ITR1 Expression | 61811 | 11 |
| YEL031W | SPF1 | Sensitivity to Pichia Farinosa killer toxin | 135259.6 | 10 |
| YOR002W | ALG6 | Asparagine-Linked Glycosylation | 62798.3 | 10 |
| YFL025C | BST1 | Bypass of Sec Thirteen | 117747 | 8 |
| YGL160W | AIM14 | Altered Inheritance rate of Mitochondria | 65849.6 | 7 |
| YGL167C | PMR1 | Plasma Membrane ATPase Related | 104554.8 | 7 |
| YLR130C | ZRT2 | Zinc-Regulated Transporter | 46342.2 | 7 |
| YBR287W |  |  | 47493.6 | 6 |
| YJR015W |  |  | 58116.5 | 6 |
| YBL011W | SCT1 | Suppressor of Choline-Transport mutants | 85693.2 | 4 |
| YKR067W | GPT2 | Glycerol-3-Phosphate acylTransferase | 83674.9 | 4 |
| YOR085W | OST3 | OligoSaccharylTransferase | 39487.6 | 4 |
| YER072W | VTC1 | Vacuolar Transporter Chaperone | 14381.3 | 3 |
| YGR089W | NNF2 |  | 106476.7 | 3 |
| Unknown subcellular location |  |  |  |  |
| YDL206W |  |  | 85973.1 | 12 |
| YOL103W | ITR2 | myo-Inositol TRansporter | 66715.8 | 12 |
| YLR241W | CSC1 | Calcium permeable Stress-gated cation Channel | 89348.1 | 11 |
| YOL158C | ENB1 | ENteroBactin | 66989.4 | 11 |
| YER060W | FCY21 | FluoroCYtosine resistance | 58041.4 | 10 |
| YHR078W |  |  | 63331.2 | 8 |
| YGR199W | PMT6 | Protein O-MannosylTransferase | 88018.7 | 8 |
| YGR149W | GPC1 | GlyceroPhosphoCholine acyltransferase | 51652.9 | 7 |
| YLR214W | FRE1 | Ferric REDuctase | 78879.7 | 7 |
| YLR443W | ECM7 | ExtraCellular Mutant | 50314.2 | 4 |
| YOR301W | RAX1 | Revert to Axial | 50160.1 | 3 |

**Table S5.**
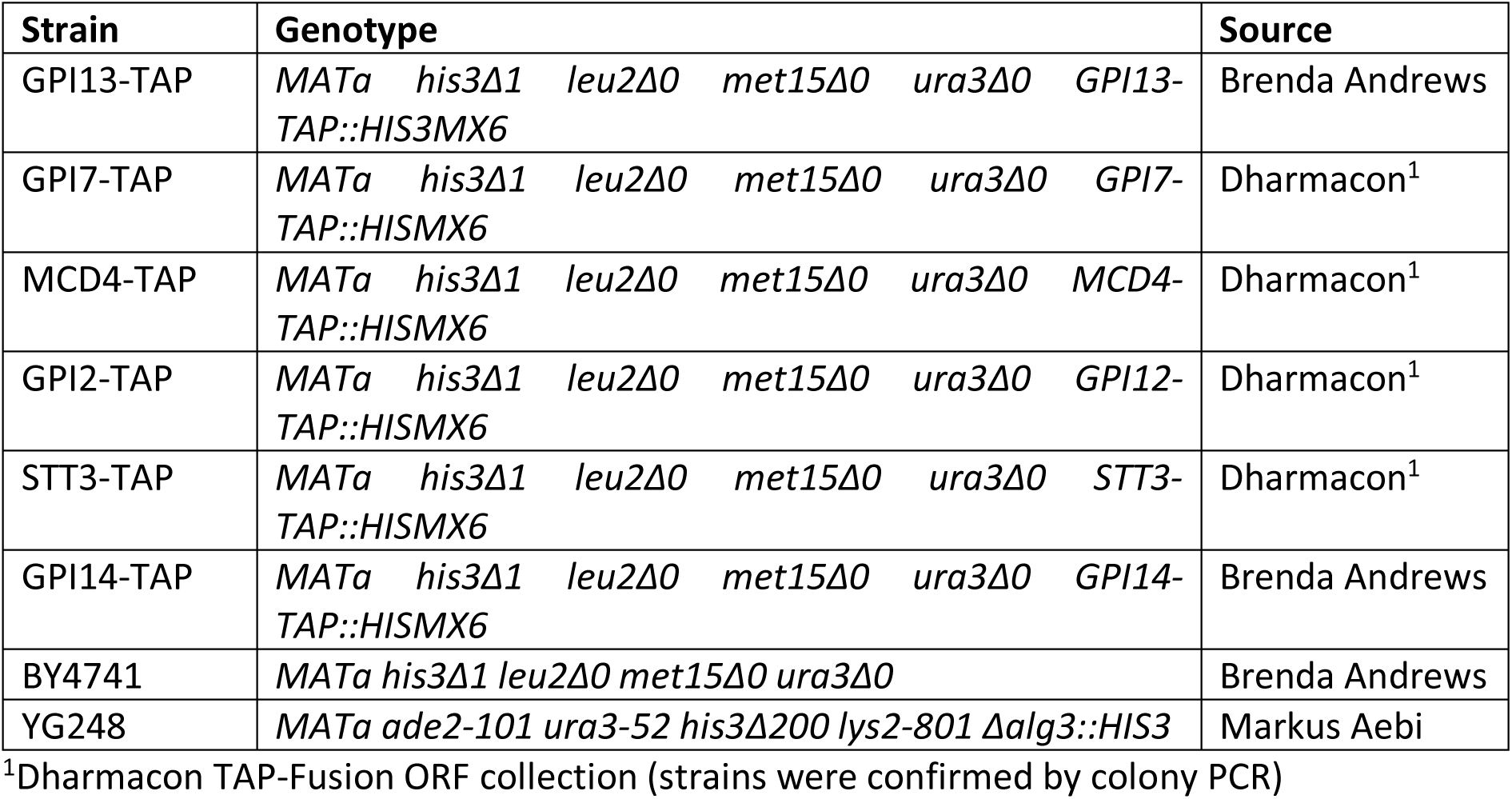
Yeast strains used in this study

**Table S6.**
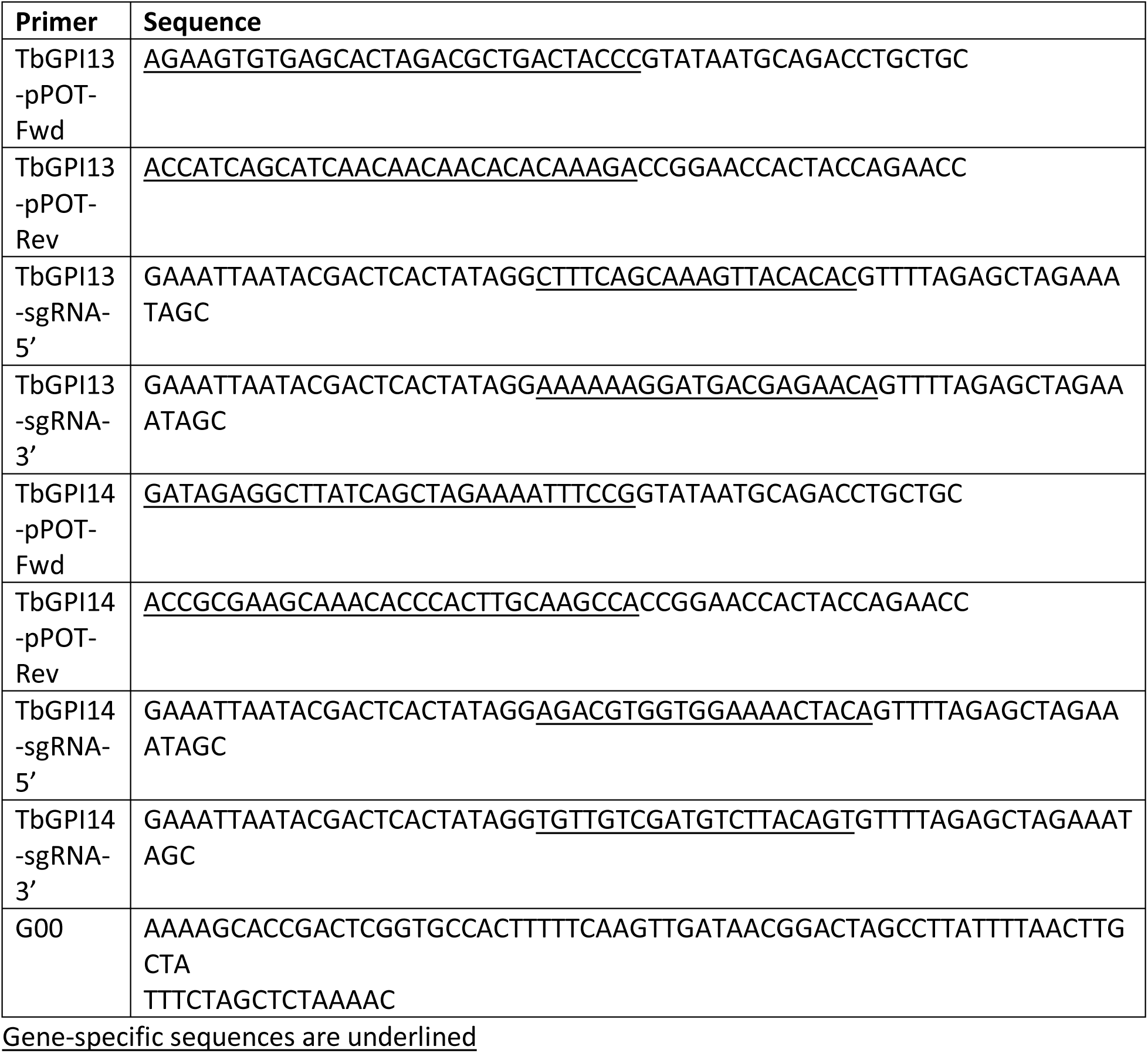
Primers used to create knockout *T. brucei* cell lines

